# Dynamic Landscapes of tRNA Transcriptomes and Translatomes in Diverse Mouse Tissues

**DOI:** 10.1101/2022.04.27.489644

**Authors:** Peng Yu, Siting Zhou, Yan Gao, Yu Liang, Wenbin Guo, Dan Ohtan Wang, Shuaiwen Ding, Shuibin Lin, Jinkai Wang, Yixian Cun

## Abstract

Although the function of tRNA in translational process is well established, it remains controversial whether tRNA abundance is tightly associated with **translational efficiency** (TE) in mammals. For example, how critically the expression of tRNAs contributes to the establishment of **tissue-specific** proteomes in mammals has not been well addressed. Here, we measured both **tRNA expression** using DM-tRNA-seq and ribosome-associated mRNAs in the brain, heart, and testis of RiboTag mice. Remarkable variation in the expression of tRNA isodecoders was observed among the different tissues. When the statistical effect of isodecoder-grouping on reducing variations is considered through permutating the anticodons, we observed an expected reduction in the tissue-variations of anticodon expression, an unexpected smaller variation of **anticodon usage bias**, and an unexpected larger variation of tRNA isotype expression. Regardless whether or not they share the same anticodons, isotypes encoding the same amino acids are co-expressed across different tissues. Based on the tRNA expression and TE computed from RiboTag-seq, we find that the tRNA adaptation index (tAI) values and TE are significantly correlated in the same tissues but not among tissues; tRNAs and the **amino acid compositions** of translating peptides are positively correlated in the same tissues but not between tissues. We therefore hypothesize that the tissue-specific expression of tRNAs might be related to post-transcriptional mechanisms, such as aminoacylation, modification, and tRNA-derived small RNAs (tsRNAs). This study provides a resource for tRNA and translation studies to gain novel insights into the dynamics of tRNAs and their role in translational regulation.

## Introduction

The genetic information is transmitted from DNA to RNA, and to proteins. However, the correlation between mRNA abundance and protein expression level is far from linear, suggesting that translation process plays an indispensable role in determining the output of proteins [1]. During protein synthesis, tRNAs decode the template mRNAs via codon-anticodon pairing and deliver the amino acids to the corresponding polypeptide chain in the ribosomes [2]. tRNAs are small non-coding RNAs, 70–90 nucleotides in length, transcribed by RNA polymerase III (RNAPIII), and constitute 4–10% of the total RNAs in a cell [3]. Although there are only 20 amino acids and 64 codons, about 400 nuclear-derived tRNAs have been annotated in mammals (*e.g.*, 429 and 401 annotated tRNA genes in human and mouse genomes respectively) in addition to 22 mitochondrial-derived tRNAs (mt-tRNAs) [4]. tRNA transcripts that carry the same anticodons but different body sequences are termed isodecoders [5], while different tRNA species accepting the same amino acids are termed isoacceptors [3]. There are 49 and 47 isoacceptors annotated in human and mouse genomes respectively [6,7].

In bacteria and yeast, the tRNA abundance correlates well with the codon usage of highly translated genes [8–10]. In mammals, the relationship is still in debate. Several studies have shown correlation between tRNA and translation. For example, Kimberly et al. reported that the tRNA abundance is significantly correlated with the codon usage of tissue-specific and highly expressed genes [11]. Hila et al. found that tRNAs induced in proliferative cells or differentiated cells often decode codons that are enriched in mRNAs related to cell-autonomy and multicellularity [12]. Yedael et al. reported better adaptation between tissue-specific genes and their tRNA pool when compared with non-specific genes [13]. Xavier et al. reported that the tRNA pool related to the proliferative state affects translational efficiency (TE) [14]. Hamed reported that codon usage is correlated with the TE in adaptation to environmental and physiological changes [15]. However, other studies have shown that the correlation is poor. Marie et al. reported that the significant differences in synonymous codon usage between tissues is not due to translational selection [16]. Kanaya et al. reported that the ribosome genes and histone genes show no difference in codon usage, implying no translational regulation through tRNAs [17]. Thus, it is unclear how tRNA expression profiles are correlated to TE of specific transcripts.

To address this issue, quantitative tRNA expression evaluation is desirable. However, due to the stable structure and diverse post-transcriptional modifications of tRNAs which interfere with reverse transcription efficiency and adaptor ligation, it has been difficult for standard sequencing methods to detect tRNA pools efficiently and quantitatively. Most studies have utilized microarrays or Pol III chromatin immunoprecipitation followed by sequencing (ChIP-seq) to identify tRNA transcriptomes. In recent years, more high-throughput sequencing methods have been developed to measure the abundances of tRNAs [18–22]. However, none of these studies have compared the tRNA abundance with matched translatome data. Therefore, whether the dynamics of tRNA expression contribute to the establishment of tissue-specific translatomes in mammals has not been well addressed.

Although still largely elusive, the regulation of tRNA expression can be possibly mediated by transcriptional and post-transcriptional mechanisms. On the one hand, the occupancies of Pol III as considered at the isoacceptor family level were invariant in multiple mammalian tissues [7]. On the other hand, the RNA modification and structure of tRNAs can regulate the ribonucleases-catalyzed degradation of tRNAs [23–25]. It was also reported that multiple tRNAs was degraded when histidine or leucine becomes limited, suggesting the tRNA expression was also under post-transcriptional regulation [26].

In this study, to overcome the difficulty of quantification of tRNAs expression, we applied the DM-tRNA-seq method reported by Zheng et al. to evaluate the diversity of tRNA pools in three mouse tissues (brain, heart, and testis). The DM-tRNA-seq utilizes engineered demethylases AlkB to remove base methylation on tRNAs and can measure the tRNA transcriptomes efficiently and quantitatively [27]. Meanwhile, we applied RiboTag-seq to capture the ribosome-associated mRNA in the same mouse tissues [28]. We found degrees of variations of tRNA expression at the isodecoder, the isoacceptor, and the amino acid level among different mouse tissues, suggesting the dynamic expression of tRNAs. We then found that the tRNA adaptation index (tAI) values were significantly correlated with TEs intra- but not inter-tissues. Our study suggests that it is unlikely that the differential tRNA expression contributes to tissue-specific translatomes but may be resulted from post-transcriptional regulation of tRNAs.

## Results

### Dynamic expression of tRNA isodecoders among different mouse tissues

In order to systematically elucidate the tissue-specificity of tRNA expression, we obtained total RNAs from three tissues (brain, heart, and testis) of adult male *CMV-Cre*: RiboTag mice and generated tRNA libraries using DM-tRNA-seq with two biological replicates (**Figure 1A**). The RPMs (Reads Per Million mapped reads) were calculated for each tRNA annotated in genomic tRNA database GtRNAdb [4] and mitochondrial tRNA database mitotRNAdb [29] (Table S1; the bioinformatic pipeline is shown in Figure S1). As shown in Figure 1B, the biological replicates of the same tissues are highly similar to each other and clustered together, suggesting tissue-specific expression of tRNAs. We noted that the tRNA expression pattern of brain tissue is less reproducible than that of heart and testis, possibly reflecting the higher cell heterogeneity of the brain. In addition, we found mt-tRNAs accounting for 11.1% of the total detected tRNAs in testis but 64.4% in heart and 38.9% in brain, which is consistent with the order of energy demand in these tissues (Figure 1C). Since the dynamics of mt-tRNAs contents are more likely reflecting the dynamics of the number of mitochondria in the cells, we focused on the dynamics of cytosolic tRNAs (cyto-tRNAs), which may relate to the translational regulation of nuclear-derived genes.

**Figure 1.**
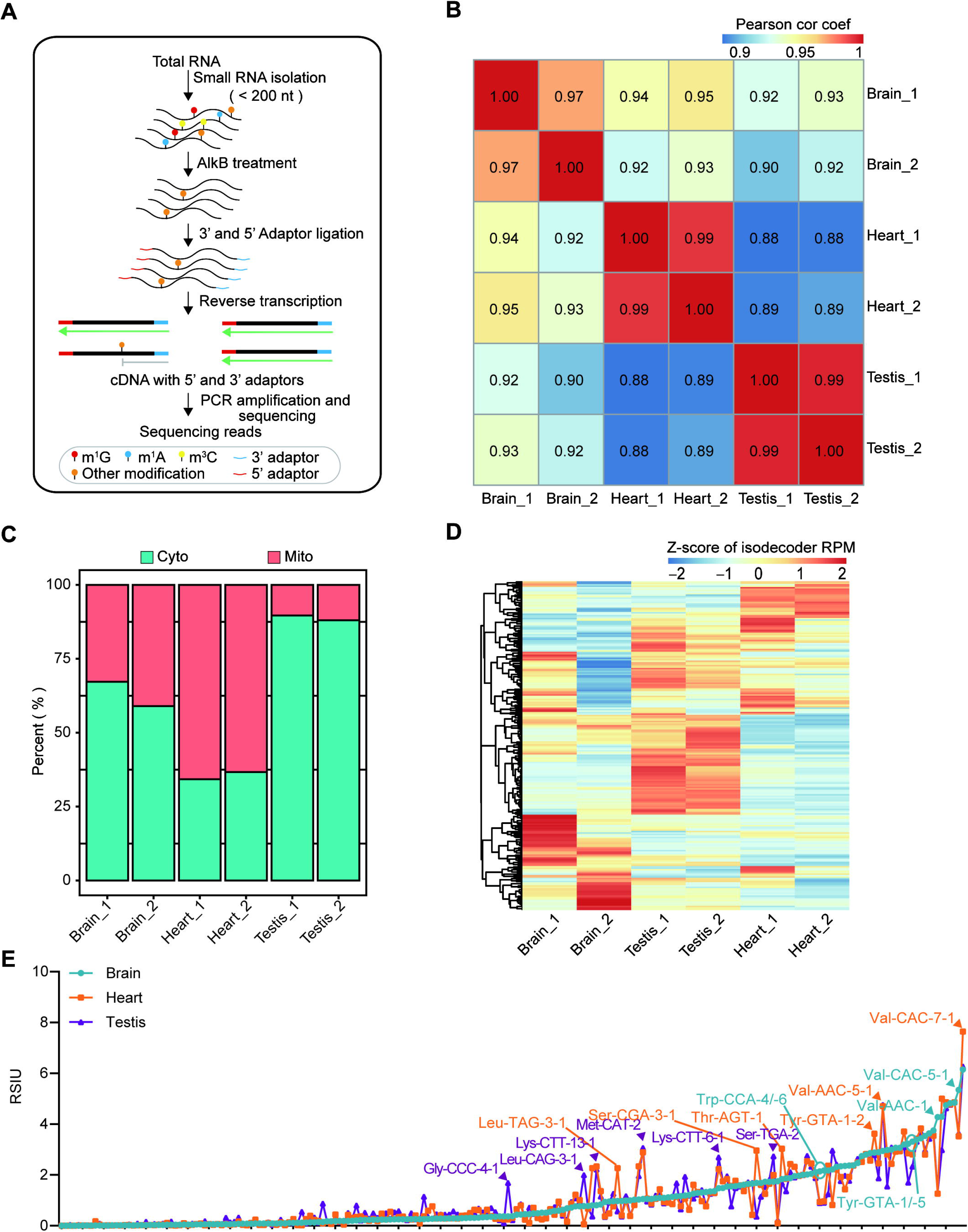
Dynamic expression of tRNA isodecoders among different mouse tissues. **A.** Schematic representation of DM-tRNA-seq based on AlkB demethylation. **B.** Heatmap of pairwise Pearson correlation coefficients of tRNA isodecoder expression among the six samples of the three mouse tissues. **C.** Stacked bar plot depicting the percentages of cytoplasmic tRNA reads and mitochondrial tRNA reads in three mouse tissues. **D.** Heatmap representing the Z-score of gene expression of tRNA isodecoders in the six samples of the three mouse tissues. **E.** Line chart comparing the strength of isodecoders usage bias in different tissues as measured by the RSIU. The representative isodecoders with high tissue specificities are indicated. Cor coef, correlation coefficient; Cyto, cytosolic; Mito, mitochondrial; RPM, Reads Per Million mapped reads; RSIU, Relative Synonymous Isodecoder Usage.

Differential expression analysis of tRNA isodecoders was performed on three tissues using DESeq2 [30]. Among the 224 detected tRNA isodecoders with unique sequences, 131 (58%) of them had significantly differential expression (FDR < 0.05) across the three tissues (Figure 1D). To further elucidate the potential role of expression regulation of tRNA isodecoders on translation, we defined a metric, Relative Synonymous Isodecoder Usage (RSIU), to analyze the usage bias of synonymous isodecoders with the same anticodons (details in ‘Materials and Methods’). As shown in Figure 1E, we observe remarkable differences in the RSIU values among the tissues. RSIU values in one of three tissues for 36 of 224 isodecoders are more than 2 folds of those in any other two tissues. For example, the RSIU value of isodecoder Gly-CCC-4-1 in testis was 6.9 and 4.5 folds of that in heart and brain, respectively.

### Tissue-specific expression of isodecoders results in tissue-specific expression but not the usage bias of anticodons

To assess tRNA expression at the anticodon level, the 224 cytosolic tRNAs identified by DM-tRNA-seq were separated into 47 groups by combining the isodecoders with the same anticodons. The expression heatmap of tRNA isoacceptors demonstrates the differential expression patterns across the three tissues (**Figure 2A**). We found the samples of the same tissues were clustered together according to the expression of the 47 anticodons, suggesting tissue-specific expression of anticodons (Figure 2A and B). Of note, we realized that the coefficient of variation (CV) of isoacceptors among tissues were significantly smaller than that of isodecoders (Figure 2C), consistent with the recent study reporting milder differences of isoacceptors among tissues [21]. To validate our results, we also compared the CV of isoacceptors and isodecoders using one published tRNA dataset examined by a different technology QuantM-tRNA seq [21]. Similarly, we observed greatly reduced CV of the expression by isoacceptors than by isodecoders (Figure S2A). To test whether the relatively smaller variation of isoacceptors expression was due to genuinely tissue-specific expression of anticodons, we made the heatmap of isoacceptors expression examined using QuantM-tRNA seq based on Z-score for each anticodon, we found reproducible tissue-specific isoacceptors expression, although different regions of brain were not largely distinct from each other (Figure S2B), which is consistent with our results, suggesting tissue-specific expression of isoacceptors.

**Figure 2.**
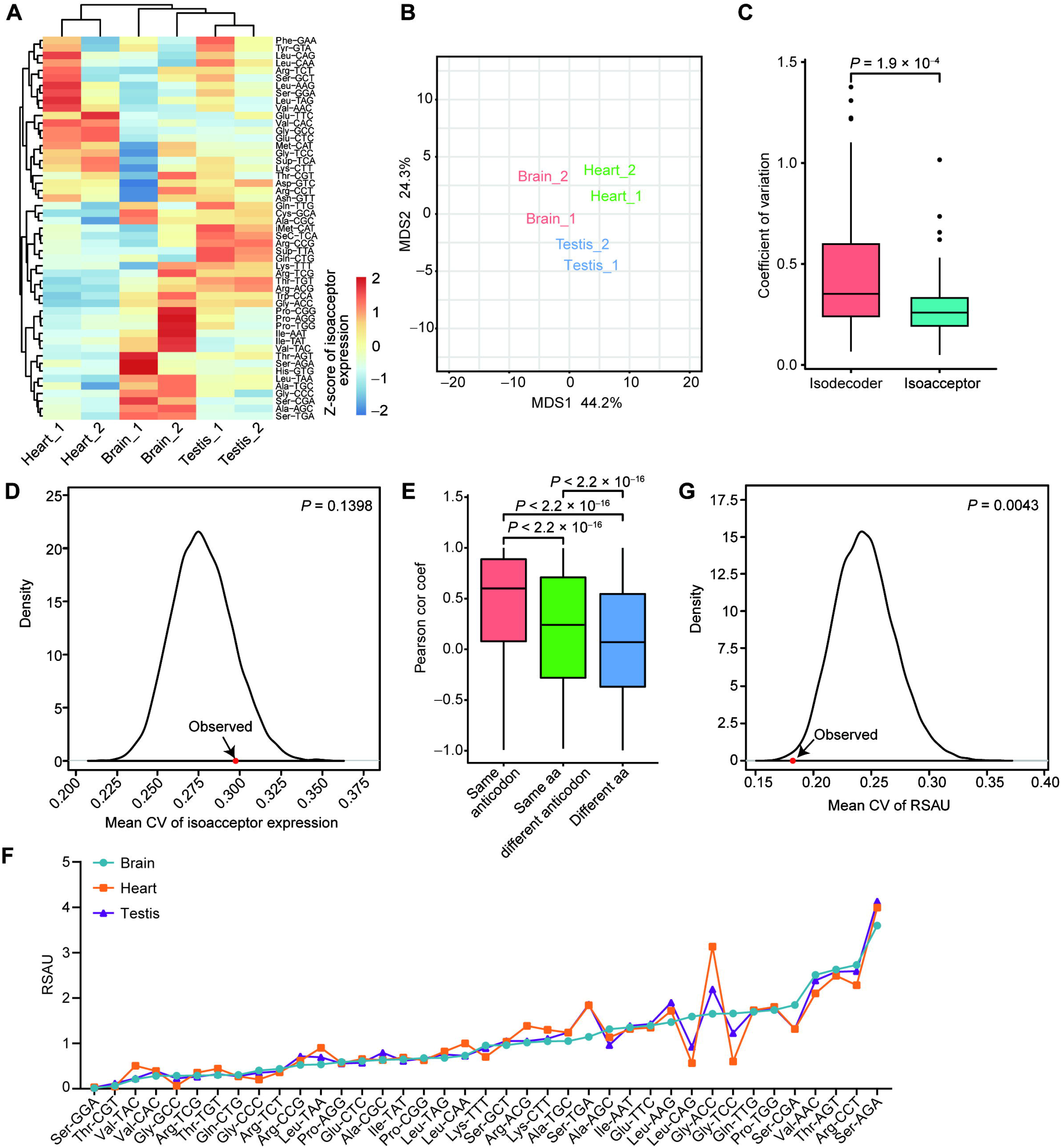
Tissue-specific expression of isodecoders result in tissue-specific expression but not the usage bias of anticodons. **A.** Heatmap representing the Z-score of tRNA isoacceptor expression in the six samples of the three mouse tissues. **B.** Multidimensional scaling plot displaying the clustering of the six samples of the three mouse tissues according to the tRNA expression profiles. **C.** Boxplot comparing the CVs of isodecoders and isoacceptors among the six samples of the three mouse tissues, *P-*values of two-tailed Wilcoxon tests are indicated. **D.** Density plot showing the distribution of mean CV of isoacceptor expression across the six samples for 10,000 permutations as well as the observed as indicated by red dot and arrow. *P-*value is calculated as the proportion of permutations with greater x-axis values than the observed. **E.** Boxplot comparing the pairwise Pearson correlation coefficients of three groups of isodecoders: “same anticodon”, “same amino acid but different anticodon”, and “different amino acid” according to the corresponding anticodons and amino acids of the pairs of two isodecoders. *P-*values of two-tailed Wilcoxon tests are indicated. **F.** Line chart comparing the strength of anticodon usage bias based on DM-tRNA-seq in three tissues as measured by the RSAU. **G.** Density plot showing the distribution of mean CV of RSAU values across the six samples for 10,000 permutations as well as the observed as indicated by red dot and arrow. *P-*value is calculated as the proportion of permutations with smaller x-axis values than the observed. CV, Coefficient of variation; aa, amino acid; RSAU, Relative Synonymous Anticodon Usage.

Then we asked why the CV of isoacceptors was smaller than isodecoders. We suspected that averaging the subgroups of isodecoders would reduce variations due to statistical principles. We therefore asked whether the reduced variations of isoacceptors among tissues were simply statistically due to the random combinations of isodecoders. For this purpose, we performed permutation analyses by randomly permutating anticodon of the isodecoders and regrouped them into isoacceptors according to the permutated anticodon. We found the CV of the observed isoacceptors among tissues was greater than 86% of 10,000 permutations, indicating a non-significant difference (Figure 2D). The results suggest that the dynamic expression of isoacceptors is simply a reflection of the dynamic expressions of isodecoders. In other words, although not so remarkable, the dynamic expression of isoacceptors is genuine.

We then turned to uncover the relationships among the expression levels of the isodecoders. We calculated the correlation coefficient of any two isodecoders across all six samples (Figure 2E). The Pearson’s correlation coefficient of isodecoder pairs encoding different amino acids, which are the most unrelated isodecoders, are around 0, suggesting the unrelated isodecoders are independently regulated. Interestingly, we found the correlation coefficients between the isodecoders pairs with the different anticodons but encoding the same amino acids were significantly greater than those isodecoders pairs encoding different amino acids. In addition, the isodecoder pairs with the same anticodons had the highest correlation coefficient. To confirm, we performed the same analyses using the published dataset of tRNA expression in multiple mouse tissues based on a different tRNA sequencing technology QuantM-tRNA seq [21]. We observed similar results that the isodecoders pairs with the same anticodons and the pairs with different anticodons but encoding the same amino acids were almost equal and both had significantly greater correlation coefficients than the unrelated pairs (Figure S2C). These results suggest that functionally related isoacceptors do not randomly fluctuate among different tissues but are associated and possibly co-regulated across different tissues, especially at the amino acid level.

Nevertheless, it is an interesting question whether the tissue-specific expressions of isoacceptors result in tissue-specific usage bias of tRNA anticodons encoding the same amino acids, which would subsequently lead to differential TEs in different tissues. We calculated the Relative Synonymous Anticodon Usage (RSAU) according to the expression of anticodons in each tissue (details in ‘Materials and Methods’). We found that the overall strength of tRNA anticodon usage bias across tissues had relatively lower diversity than tRNA isodecoder usage bias (Figure 2F and 1E). We further found that the CV of RSAU values among three tissues was significantly smaller than random permutations (Figure 2G). CV of permutated RSAU values greater than the mean of observed CV could be obtained 9957 times out of 10000 permutations, suggesting that the variations of synonymous anticodons usage bias among different tissues are prohibited. The above results demonstrate that the distinctive expression of tRNA isoacceptors does not play a vital role in selecting specific synonymous anticodons or determining the TEs in different tissues.

Because the isodecoders encoding the same amino acids tend to be co-regulated, we speculated that the diversity of tRNA pools is most likely to match the amino acid composition within specific physiological states during the translation process. We then tested whether the tRNA isotype expression at amino acid level also had tissue specificities by combing the tRNAs encoding the same amino acids. As shown in Figure 3A and B, there is an obvious tissue-specific tRNA isotype expression. In addition, the mean CV of isotype expression across these three tissues is greater than all 10,000 permutations, suggesting there is genuine tissue-specific isotype expression (Figure 3C and D). These results together with the above results imply that the dynamic regulation of tRNAs among tissues is more likely a reflection of tissue-specific needs of tRNAs encoding specific amino acids rather than optimizing the codon usages for efficient translation.

**Figure 3.**
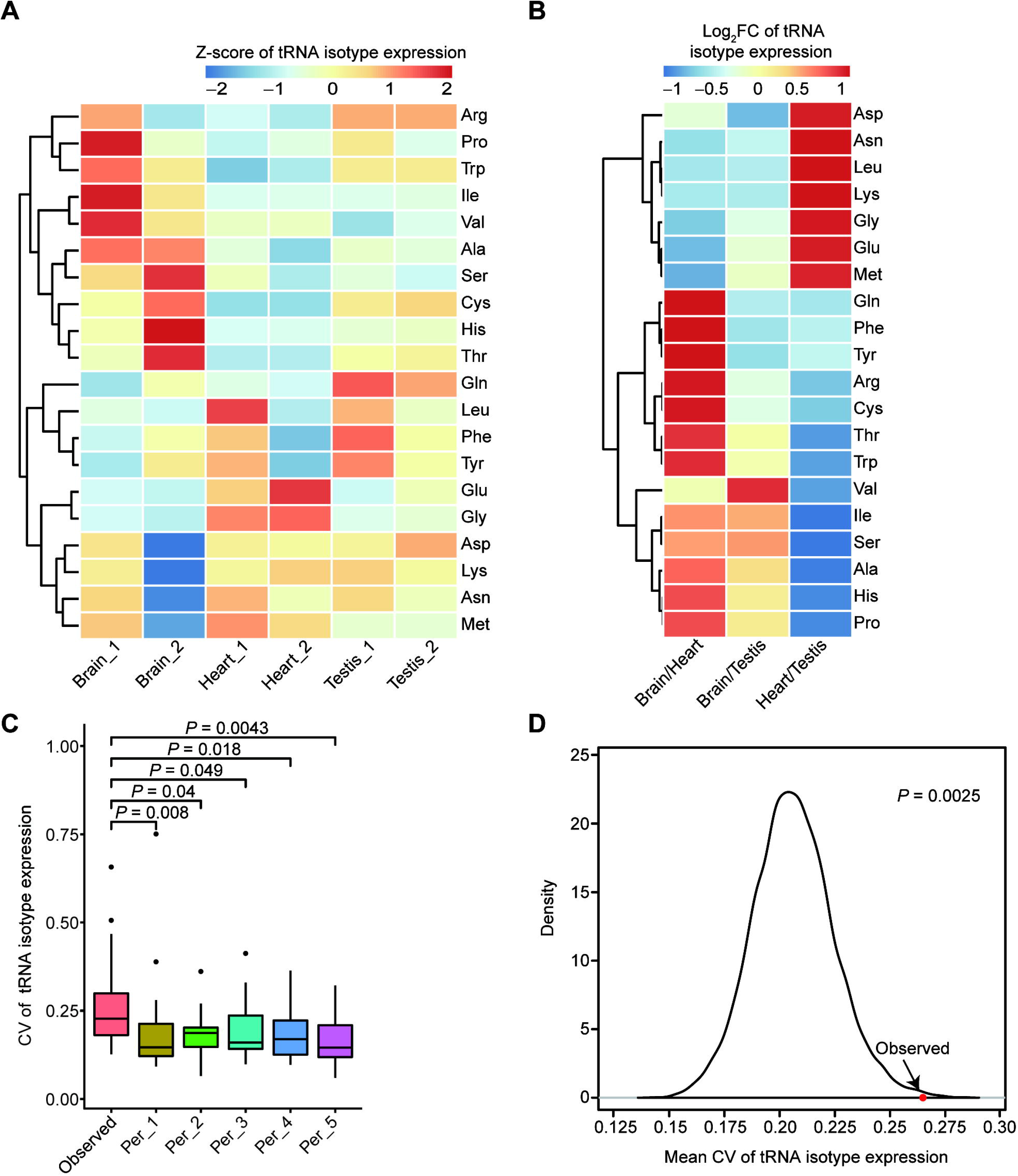
Tissue-specific tRNA isotype expression at amino acid level. **A.** Heatmap representing the Z-score of tRNA isotype expression in the six samples of three mouse tissues. **B.** Heatmap representing the log_2_-transformed fold change of tRNA isotype expression for the pairwise comparisons of the three tissues. **C.** Box plots comparing the CV of tRNA isotype expression across six samples (observed) with five random permutations, *P*-values of two-tailed Wilcoxon tests are indicated. **D.** Density plot showing the distribution of mean CV of tRNA isotype expression across the six samples for 10,000 permutations as well as the observed as indicated by red dot and arrow. *P-*value is calculated as the proportion of permutations with greater x-axis values than the observed. FC, fold change.

### RiboTag-seq analysis of translatomes in multiple mouse tissues

To further elucidate whether dynamic tRNA expression contributes to the establishment of tissue-specific translatomes, we performed RiboTag-seq in the same samples we applied DM-tRNA-seq to. The RiboTag-seq technology takes advantage of RPL22, a component of the 60S subunit of ribosome, to capture the actively translating ribosomes (**Figure 4A**). The expression of RPL22-HA protein can be activated by Cre recombinase-mediated replacement of exon 4 with an HA-tagged exon 4 of *Rpl22* gene [31]. To create a line of mice constitutively expressing Rpl22-HA protein in multiple tissues, RiboTag mice were mated with *CMV-Cre* mice (Figure 4B). We validated the heterozygote *CMV-Cre* and homozygous *Rpl22-HA* alleles in the genomes of offspring and confirmed expression of RPL22-HA protein in a plurality of tissue homogenates, followed by efficient immunoprecipitating (Figure 4C and Figure S3A–C).

**Figure 4.**
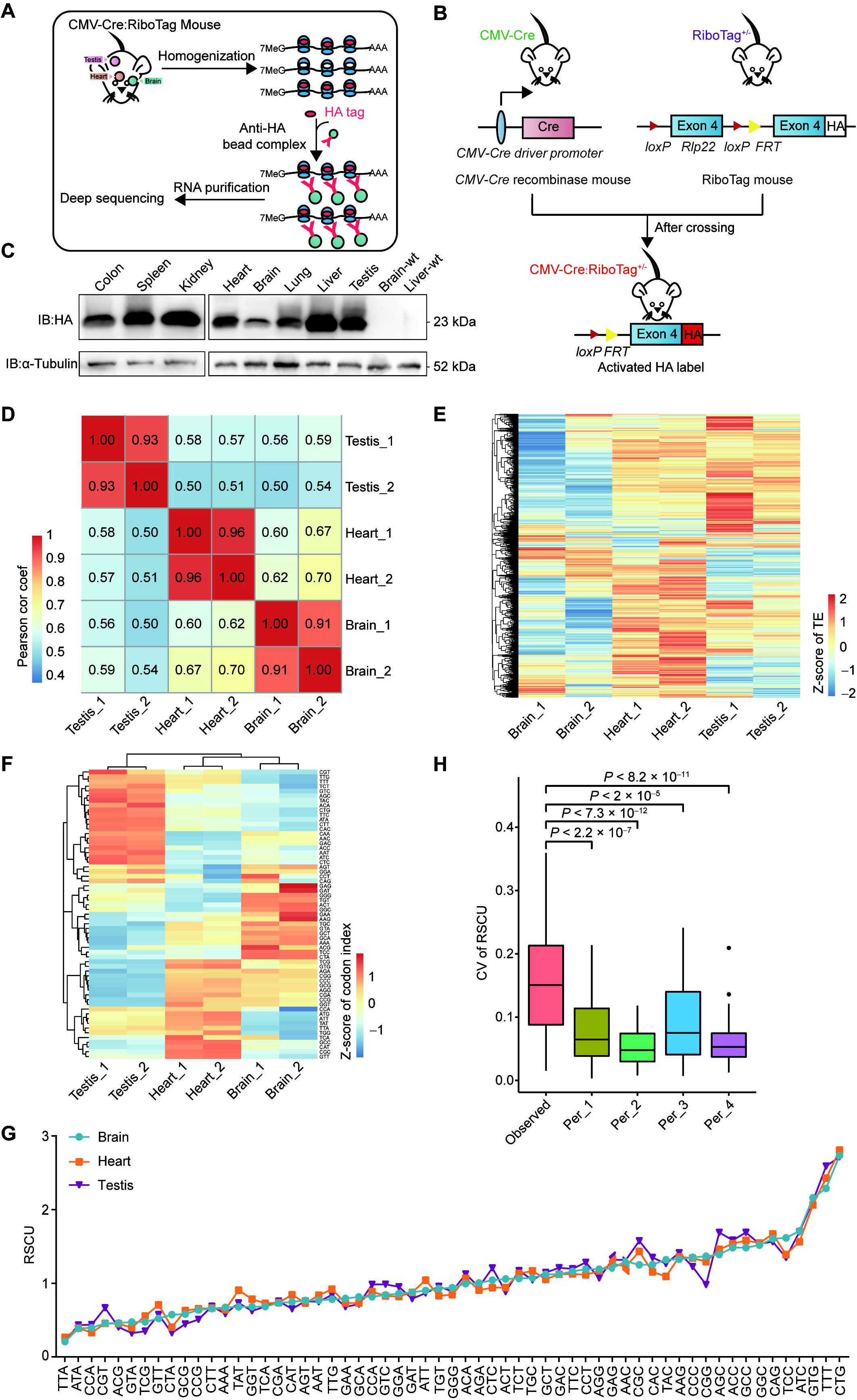
RiboTag analysis of translatomes in multiple mouse tissues. **A.** Overview of RiboTag technology. **B.** Diagram depicting the RiboTag mouse systems. **C.** Western blot analysis of RPL22-HA in different tissues of *CMV-Cre*: RiboTag mouse. **D.** Heatmap of pairwise Pearson correlation coefficients among the six mouse samples in three tissues based on the FPKMs of genes in Input samples of RiboTag. **E.** Heatmap representing the Z-score of TEs in six samples of three mouse tissues. **F.** Heatmap representing the Z-score of codon indexes of top 5% highly translated genes in six samples of three mouse tissues. **G.** Line chart comparing the strength of codon usage bias in different tissues as measured by the RSCU. **H**. Boxplots comparing the CV of RSCU values of the top 5% of highly translated genes (observed) across six samples with four random permutations, *P-*values of two-tailed Wilcoxon tests are indicated. HA, hemagglutinin; MeG, N7-methylated guanosine; wt, wild type; FPKM, Fragments Per Kilobase per Million mapped of fragments; TE, translational efficiency; RSCU, Relative Synonymous Codon Usage.

Since RiboTag only sequenced the RNAs bound by the translation factor RPL22, we calculated the translation levels, which were represented by the gene expression levels of immunoprecipitation RNAs (IP), as well as TEs, which were the translation levels normalized by the expression of input RNAs (Table S2; the bioinformatic pipeline is shown in Figure S1). Strong tissue-specific gene expression as well as TEs were observed (Figure 4D and E). Gene ontology (GO) analysis and Kyoto encyclopedia of genes and genomes (KEGG) analysis of the highly translated gene (top 5%) revealed enrichment for tissue development or tissue physiology-related processes and pathways (Figure S4A–C). We then asked whether the composition of codons was different among these tissues. For this purpose, we defined a metric of codon index (details in ‘Materials and Methods’), which is the proportion of specific codons in all of the top 5% highly translated genes weighted by the translation level of each gene. As shown in Figure 4F, there are distinctive codon indexes among the three tissues, suggesting it might be necessary for tissue-specific tRNA pools. To test whether the usage biases of synonymous codons also differ among tissues, we utilized the previously defined metric Relative Synonymous Codon Usage (RSCU) for each codon [32], which is the observed frequency of specific codons divided by the frequency expected under the assumption of equal usage of the synonymous codons. In the top 5% highly translated genes of each tissue, we found moderate differences among brain, heart, and testis (Figure 4G). However, the CV of RSCU values among the three tissues based on highly translated genes was still significantly higher than the CV based on randomly sampled 5% genes, suggesting that codon usage bias of highly translated genes is truly differential among tissues (Figure 4H).

### tRNA pools adapt better to highly translated genes in the same tissues but not to tissue-specifically translated genes

The more accurate interaction analysis between mRNAs and cognate tRNAs will provide a pivotal way for evaluating effective and accurate translation [33,34]. To further elucidate the intrinsic relationship between tRNA expression and mRNA translation, we integrated the data of tissue-specific DM-tRNA-seq and RiboTag-seq to comprehensively uncover the correlation between tRNA pools and codon usage bias in highly translated genes. The adaptation of a specific gene to a specific tRNA pool in terms of codon usage bias can be well evaluated using a widely used metric tRNA adaptation index (tAI) [35,36]. We found the highly translated genes had significantly higher tAI values than moderately and lowly translated genes in all the three tissues based on the tRNA pools of the corresponding tissues (**Figure 5A**), suggesting the role of tRNA in regulating translation in certain tissues. However, when we tested the adaption of highly translated genes with the tRNA pools from other tissues, we found the highly translated genes did not show the highest tAI values based on the tRNA pool of the same tissues. Instead, the tRNA pool of heart had the best adaptation with the highly translated genes of all tissues (Figure 5B). In addition, we performed the correlation analysis between isoacceptor abundances and the codon compositions of the top 5% highly translated genes of each tissue based on the general codon-anticodon recognition rules for tRNA genes [36]. Similar to the tAI analyses, we found significant correlations in heart and testis but not between tissues (Figure S5A and B). The above results suggest that although highly translated genes require tRNAs, the tissue-specific regulation of tRNA expression is not intended to better adapt the tissue-specific usage bias of synonymous codons. In other words, mammals are not likely to regulate tissue-specific translation of certain genes through regulating the composition of tRNA pools, which is consistent with the observation that the usage biases of anticodon do not show significant differences among different tissues (Figure 2G).

**Figure 5.**
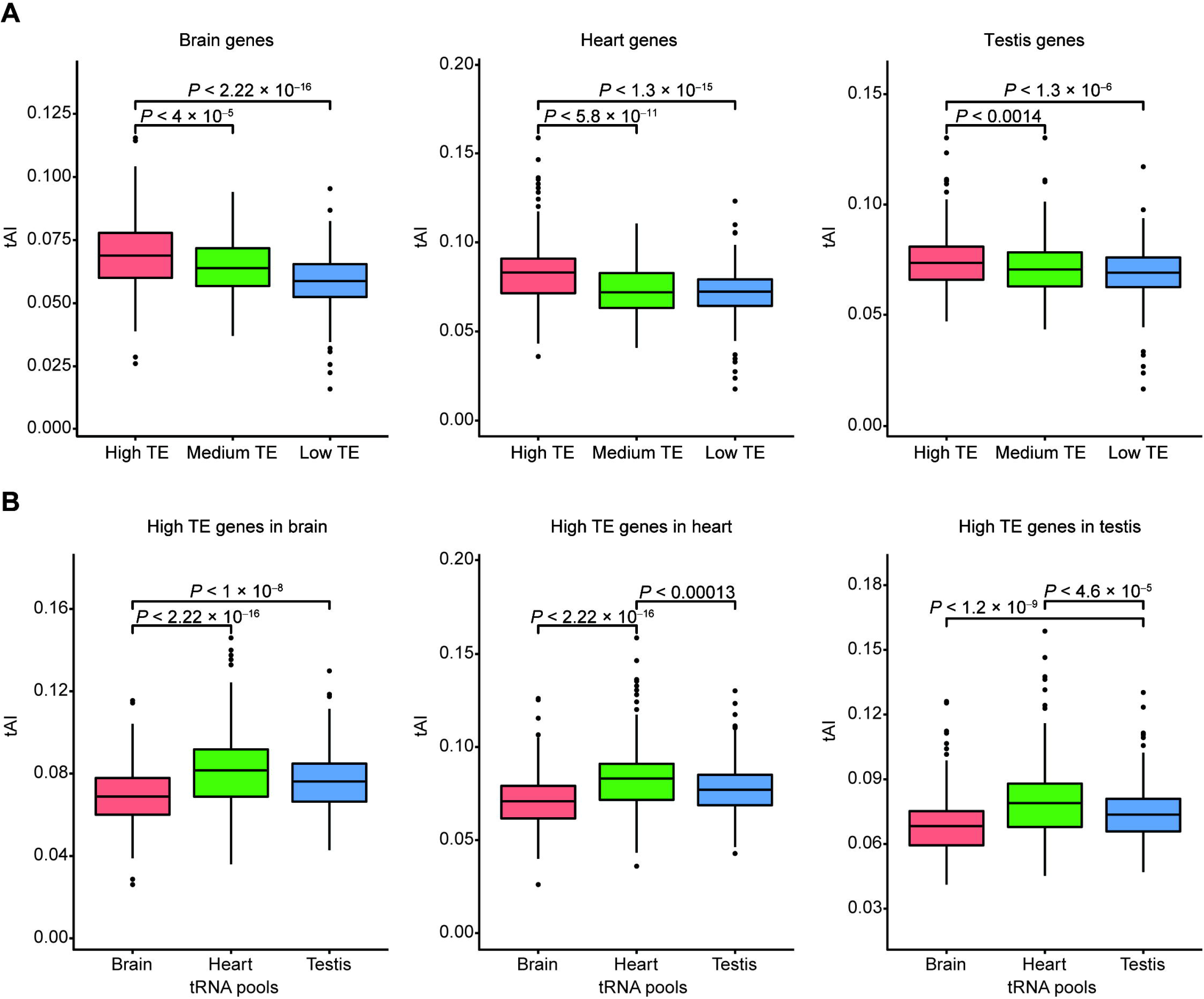
tRNA pools adapt better to highly translated genes in the same tissues but not to tissue-specifically translated genes. **A.** Box plots comparing the tAI of genes with different TE levels based on the tRNA pools of the same tissues in the three tissues respectively. *P-*values of the two-tailed Wilcoxon tests are indicated. **B.** Box plots showing the tAI of the top 5% of highly translated genes in the brain (left panel), heart (middle panel), and testis (right panel) calculated based on the tRNA pools of the three tissues respectively. *P-*values of two-tailed Wilcoxon tests are indicated. tAI, tRNA adaptation index.

### tRNA expression correlates with amino acids composition in the same tissues but not between tissues

The analyses of tRNA expression across diverse tissues revealed that isodecoders encoding the same amino acids are likely co-regulated, suggesting that the dynamics of tRNA expression in different tissues might be related with different amino acids compositions of peptides in different tissues. To test this hypothesis, we first tested whether the amino acids compositions are different across the translatomes of different tissues. We calculated each amino acid composition by summing up the number of codons encoding the amino acid of the top 5% highly translated genes weighted by the translation level (RPKM of IP). As shown in **Figure 6A**, we observed reproducible tissue-specific amino acids compositions, which is consistent with our observations that the tRNA isotype expression is tissue-specific (Figure 3A). We also found a positive correlations between the amino acid compositions and the tRNA isotype expression in heart (P = 0.023), and a trend of positive correlation in brain (P = 0.067) and testis (P = 0.11) respectively (Figure 6B). To further address whether the tissue-specific tRNA expression is related to the tissue-specific amino acid composition of peptides, we tested the correlation between amino acid compositions subtracted by the means and Z-score of tRNA isotype expression among the three tissues. We observed no significant correlation between them (Figure 6C). A non-significant correlation was also observed when we compared the differences of amino acid composition and the differences of tRNA isotype expression between any two tissues (Figure S6A and B).

**Figure 6.**
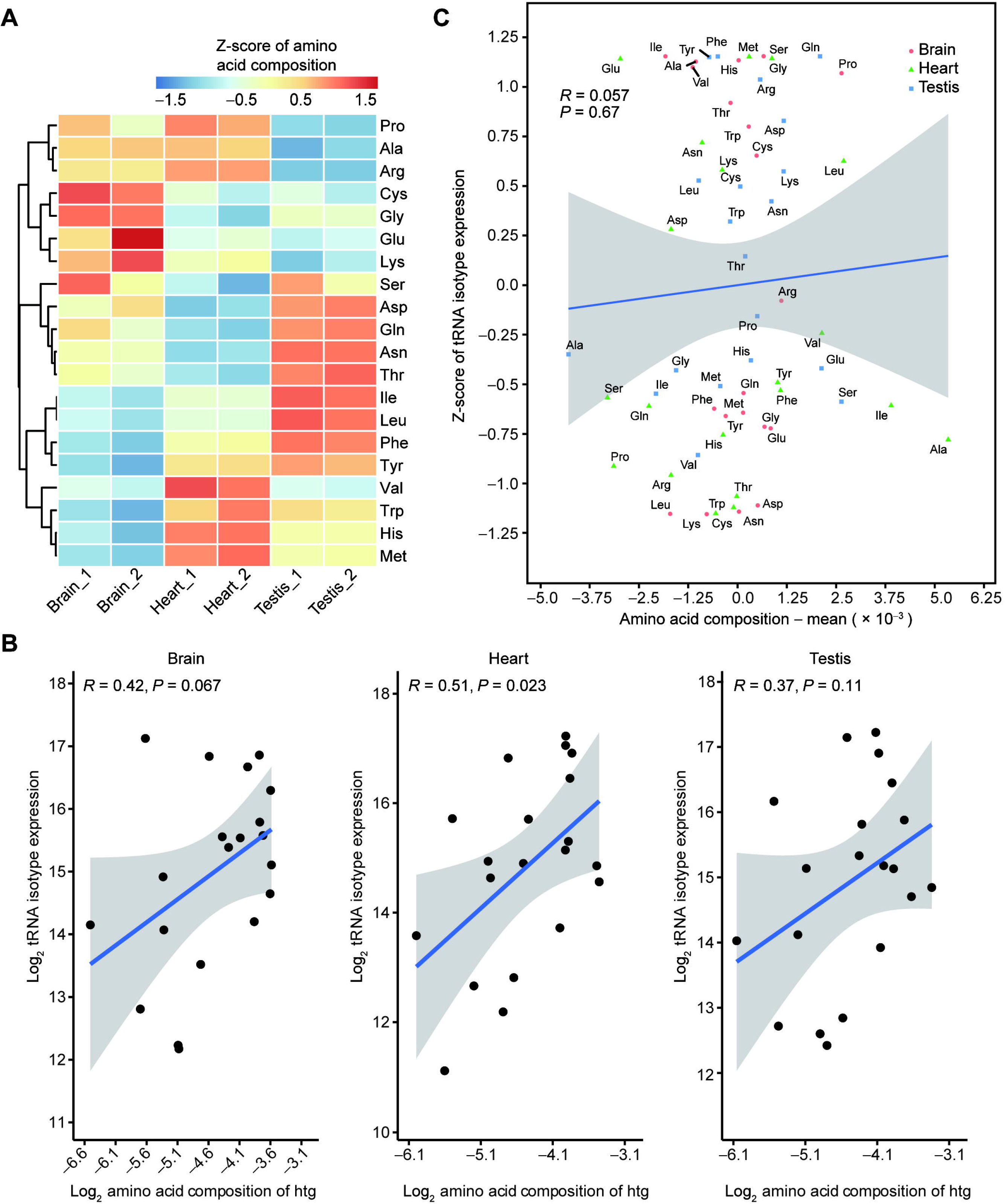
tRNA expression correlates with amino acids composition in the same tissues but not between tissues. **A.** Heatmap representing the Z-score of amino acid compositions of top 5% highly translated genes in the six samples of three mouse tissues. **B.** Scatter plots showing the correlation of amino acid compositions of the top 5% highly translated genes with the tRNA isotype expression in brain (left panel), heart (middle panel), and testis (right panel). Blue lines indicate fitted linear models and Pearson’s correlations are shown. **C.** Scatter plots showing no significant linear correlation of amino acid compositions subtracted by means with the Z-score of tRNA isotype expression in all three tissues. htg, highly translated genes.

We found that the isodecoders encoding the same amino acid were co-regulated across different tissues (Figure 2E). Based on the above results, this co-regulation is not likely due to active regulatory mechanisms to control the translatomes in a tissue-specific manner. On the contrary, it might be due to post-transcriptional regulation of tRNAs, such as tRNA modification and aminoacylation, the attachment of amino acids to tRNA.

## Discussion

Although it is well known that tRNAs play a vital role in the synthesis of protein, whether the tRNA pool correlates well with translational efficiency is obscure. Here, based on multiple measurements of tRNAs and translatomes in multiple mouse tissues, we confirmed genuinely dynamic expression of tRNA isodecoder pools as well as isoacceptors among three mouse tissues. Meanwhile, the tRNA pool is significantly correlated with translational efficiency and amino acid composition of the highly translated gene in the same tissues but not between tissues. We finally propose that the tissue-specific expression of tRNA may be due to post-transcriptional regulation.

Interestingly, tRNA expression is significantly correlated with translational efficiency in the same tissues but not between different tissues. Consistently, several studies have reported that tRNA-codon bias co-adaptation is not tissue-specific but globally driven [13,37]. These results together suggest the organisms may not regulate the translation of specific genes tissue-specifically through regulating tRNA expression, probably due to the difficulty of achieving precise adjustment through the regulation of tRNA expression. Nevertheless, we cannot rule out that there may be a weak correlation to be revealed and more accurate detection methods need to be developed in the future.

It has advantages of using RiboTag to measure the translational efficiency in this study. In contrast to ribosome profiling (Ribo-seq), which measures the translation through obtaining the mRNA fragments protected by ribosomes [38], the RiboTag takes advantage of RPL22, a component of the 60S subunit of the ribosome, to pull down the mRNAs involved in translation elongation. In principle, Ribo-seq has difficulty in distinguishing the large and small ribosome subunits, and thus cannot distinguish translation initiation and elongation. In contrast, RiboTag captures full-length mRNAs bound by actively translating polysomes, thus providing a more specific measurement of translation elongation. Since translation initiation and elongation may relate to translational efficiency in different manners [39], RiboTag overcomes the drawback of ribosome profiling. In addition, considering we have found a significant correlation between tAI and translational efficiency, the RiboTag technology used in this study is reliable in representing the translatome [28].

In this study, we hypothesize that the difference of tRNA between tissues is due to passive post-transcriptional regulation during the process of tRNA maturation. First, we found that the isodecoders encoding the same amino acid are co-regulated. Second, there is no difference of the Pol III binding on tRNA genes at the isoacceptor level among tissues [7], suggesting that differences in tRNA may be related to post-transcriptional regulation. In addition, it was reported that in *Escherichia coli*, tRNA can be destabilized and degraded in the case of amino acid starvation and upon the demand for protein synthesis decreases, suggesting the content of tRNA is related to the concentration of the free amino acids [26]. Meanwhile, several groups have shown that certain amino acids such as Cysteine [40], Glycine [41], Serine [42], and Threonyl [43] have key impacts on the modifications of tRNAs, and some modifications of tRNA will further affect tRNA abundances [44,45]. Therefore, post-transcriptional regulation of tRNA may also contribute to the tissue-specific expression of tRNAs and translatomes. This manner of tRNA regulation passively fine-tune the tRNA expression in a tissue-specifical manner but not for the purpose of regulating the translatomes.

One possible post-transcriptional regulation that may result in tRNA differences between tissues is through the aminoacylation process, which might be regulated by free amino acids concentration and the activity of aminoacyl tRNA synthetases. The activities of aminoacyl-tRNA synthetases (aaRS) are dynamic [46]. Mammals have twenty cytosolic aaRSs, which are the enzymes that attach amino acids to tRNAs and thus allow tRNA molecules to act as adaptors to decode mRNA. Individual tRNA isotype is aminoacylated by a specific aaRS. The aminoacylated tRNA is captured by a translation elongation factor and it is delivered to the ribosome for protein synthesis. The expression of tRNA isotype and free amino acids concentration may affect the level of aminoacyl-tRNAs, which in turn may have positive or negative feedback on the early processing steps of tRNAs or affect the stability of tRNAs in a tissue-specific manner, thus leading to the observed dynamic expression of tRNAs.

Another post-transcriptional regulation that may result in tRNA differences between tissues is tRNA modification. tRNAs are the most generally modified RNA species in cells. Eukaryotic tRNAs contain an average of 13 modified bases per molecule. Modifications occurring in the anticodon loop are essential to regulate mRNA decoding, while modifications outside of the anticodon loop are vital to regulate tRNA stability, tRNA localization, and tRNA folding [23]. Dynamic variations at the level of tRNA modification play a role in regulating the translational efficiency and accuracy of particular genes that rely on the codon usage. However, the profiling of tissue-specific tRNA modification is still lacking. In the future, the development of novel large-scale methods to reveal the tRNA modification level can point the light way to understand the diverse function of tRNAs during translation process.

Since it is known that DM-tRNA-seq can also generate a large fraction of incomplete tRNA reads due to the incomplete erasure of the modifications on tRNAs [22], the difference of tRNA read length also reflects the differences of modifications. According to the percent of reads with length > 40 bp, we found the proportions are quite similar between different tissues but the proportion of mt-tRNAs is larger than cytosolic tRNAs (Figure S7). This result is consistent with the previous report that cytosolic tRNAs and mitochondrial tRNA are modified differently. mt-tRNAs of higher eukaryotes have smaller and shorter stem and loop regions than that of cyto-tRNAs [23]. Modifications in mt-tRNAs are less diverse comparing with cyto-tRNAs [41,47]. m^1^A9 and m^2^G10 are considerably abundant modifications identified in mt-tRNA species [41], which can be removed by AlkB demethylases [27] and result in longer mt-tRNAs reads in DM-tRNA-seq.

In addition, tRNA modifications also contribute to different biogenesis of tRNA-derived small RNAs (tsRNAs), which are known to regulate translation in versatile ways [48]. Based on the expression of tsRNAs in brain, heart, and testis examined by the PANDORA-seq [49] and CPA-seq [50], we found the expression of tsRNAs was significantly and positively correlated with the expression of tRNAs in the same tissues and between different tissues (Figure S8A and B). The results suggest that tissue-specific expression of tRNA might be related to tsRNAs. It is possible that there might be unknown mechanisms that dynamically regulate the expression of tRNAs in different tissues in order to dynamically generate tsRNA in different tissues.

## Materials and Methods

### Animals

Mice were maintained on a 12 h light/dark cycle. The RiboTag mice (Stock No. 011029, Jackson Laboratory, Bar Harbor, Maine) and *CMV-Cre* mice (Stock No. 006054, Jackson Laboratory) were purchased from Jackson Laboratory. The RiboTag mice were bred to the *CMV-Cre* mice to obtain homozygous mice constitutively expressing *Rpl22-HA*. Once the model of *Rpl22-HA*-expressing homozygous mice was built successfully, we maintained the colony as a separate mouse line.

### Tissue sample preparation and RNA isolation

All mouse tissue samples were isolated from adult male *CMV-Cre*: RiboTag mice using procedures approved by the Animal Research Committee of Sun Yat-sen University, the First Affiliated Hospital. Samples were rapidly frozen in liquid nitrogen and stored at −80°C until use. 1 ml of TRIzol (Catalog No. 15596026, Invitrogen, Carlsbad, CA) was added per 100 mg of dissected whole tissue and samples were homogenized in TRIzol buffer with a homogenizer (JX-2010, China) until the suspension was completely homogeneous. Cell debris was removed by a high-speed centrifugation procedure. RNA was isolated according to the manufacturer’s instructions of TRIzol reagent and resuspended in nuclease-free water and stored at −80°C until DM-tRNA-seq.

### Recombinant Protein Purification

Recombinant wild-type and D135S AlkB proteins were purified as previously described [51]. pET30a-AlkB and pET30a-AlkB-D135S were transformed into BL21 bacteria for induced expression of recombinant proteins. Bacteria were inoculated and cultured LB medium at 37°C. Recombinant wild-type and D135S AlkB protein expressions were induced in BL21 bacteria (OD 0.6–0.7) using 0.5 mM IPTG (Catalog No. I5502, Sigma, St. Louis, MO) at 20°C overnight. Then the bacteria were collected and lysed by sonication, centrifuged at 15,000 rpm at 4°C for 60 min. The supernatant was collected for the purification of recombinant proteins using Ni-NTA Agarose (Catalog No. 30210, Qiagen, Alameda, CA) following the manufacturer’s instructions and stored at −80°C.

### DM-tRNA-seq

DM-tRNA-seq was performed following the previously reported protocol [27,47] with some modifications. Small RNAs (< 200 nt) were first purified using the Quick-RNA Microprep kit (Catalog No. R1050, Zymo Research, Orange, CA). Isolated small RNAs were treated with recombinant wild-type and D135S AlkB proteins to remove the dominant methylations on RNAs. Then demethylated RNAs were purified with Oligo Clean & Concentrator kit (Catalog No. D4060, Zymo Research). After that, AlkB-treated RNA libraries were constructed with NEBNext Small RNA Library Prep Set (Catalog No. E7330S, New England Biolabs Inc., Ipswich, MA). The cDNA libraries were sequenced on Illumina Hiseq X10 with paired-end 2×150 bp read length.

### Western Blotting

Tissue-specific lysates were extracted with RIPA buffer by a homogenizer. Western Blot assays were performed as described previously [52]. Nitrocellulose membranes were blocked using 5% Blotting Grade Blocker Non-Fat Dry Milk (Catalog No. 1706404XTU, Bio-Rad, Hercules, CA) and were then incubated with primary antibody at 4°C overnight. For primary antibodies, anti-HA tag (Catalog No. ab9110, Abcam, Cambridge, UK) was purchased from Abcam; anti-IgG (Catalog No. B900620, Proteintech, China), anti-tubulin (Catalog No. 11224-1-AP, Proteintech) were purchased from Proteintech. The blots were then incubated with horseradish peroxidase-conjugated secondary antibody (Catalog No. 7074, Cell Signaling Technology, Berkeley, CA) at room temperature for 1 h, and the proteins were then detected using the ECL chemiluminescence system (Tanon 4600, China).

### Polysome immunoprecipitation

RiboTag immunoprecipitation was performed as previously described [31] with some modifications. Tissue samples were extracted from *CMV-Cre*: RiboTag mice, flash-frozen in liquid nitrogen, and stored at −80°C until use. Tissues were homogenized in ice-cold homogenization buffer (50 mM Tris, pH 7.4, 1% NP-40, 100 mM KCl, 12 mM MgCl_2_, 100 μg/ml cycloheximide (Catalog No. 66819, Sigma), 1:100 protease inhibitors cocktail (Catalog No. 4693116001, Roche, Mannheim, Germany), 1 mg/ml Heparin, 1 mM DTT, 200 units/ml RNasin (Catalog No. N2111, Promega, Madison, WI) in RNase free DDW) with a homogenizer until the suspension was completely homogeneous. To remove cell debris, the homogenate was transferred to a microcentrifuge tube and centrifuged at 13,000*g* at 4°C for 15 min. Supernatants were transferred to a fresh microcentrifuge tube on ice, and then 70 μl was removed for input fraction analysis and 8 μl (8 µg) of anti-HA antibody (Catalog No. ab9110, Abcam) was added to the supernatant, followed by 4 h of incubation with slow rotation in a cold room at 4°C. Meanwhile, Pierce^TM^ Protein A/G Magnetic Beads (Catalog No. 88803, Thermo Fisher Scientific, Waltham, MA), 80 μl per sample, were equilibrated to homogenization buffer by washing three times. At the end of 4 h of incubation with antibody, beads were added to each sample, followed by incubation overnight at 4°C. The following day, samples were placed in a magnet on ice, and supernatants were recovered before washing the pellets three times for 10 min in high salt buffer (50 mM Tris, pH 7.4, 1% NP-40, 300 mM KCl, 12 mM MgCl_2_, 100 μg/ml cycloheximide, 1 mM DTT). At the end of the washes, beads were magnetized and excess buffer was removed. To prepare total RNA, 5 volumes of Qiagen RLT buffer were added to the remaining pellets or the input samples. Total RNA was prepared according to the manufacturer’s instructions using RNeasy Mini kit (Catalog No. 74104, Qiagen) and quantified with a NanoDrop 3000 spectrophotometer (Thermo Fisher Scientific) and taken for RNA-seq. For high-throughput sequencing, both input and IP samples were used for library construction with the SMARTer Stranded Total RNA-seq Kit v2 (Catalog No. 635005, Takara, Dalian, China), and single-end 50 bases reads were generated on the BGIseq500 platform (BGI-Shenzhen, China).

### Processing of high-throughput sequencing data

The nuclear and mitochondrial tRNAs reference sequences were downloaded from GtRNAdb [4] and mitochondrial tRNA database mitotRNAdb [29], respectively. Nuclear and mitochondrial tRNAs with unique sequences generated by collapsing the identical tRNAs were merged and used as the reference for downstream mapping. DM-tRNA-seq raw reads were first processed using Cutadapt v1.18 to remove adaptor sequences and 3′-CCA sequences and to discard reads shorter than 25 nt. Then, Bowtie2 (v2.3.5) [53] was used to align the adaptor-trimmed and filtered reads to the tRNA reference sequences of the mouse genome (mm10) with the parameters: --min-score G,1,8 --local -D 20 -R 3 -N 1 -L 10 -I S,1,0.5. Only reads with unique hits and mapping quality > 10 were considered for further analysis. The isodecoder RPMs (Reads Per Million mapped reads) were calculated by multiplying the number of reads mapped to the gene by 10^6^ and dividing it by the total number of mapped reads. The anticodon-level or amino acid-level counts were calculated by summing up the counts of isodecoders with the same anticodons or encoding the same amino acids. tRNA-seq read count tables at both the anticodon-level and isodecoder-level were used to perform differential tRNA expression analysis between each two of the three mouse tissues using the DESeq2 [30]. Differentially expressed tRNAs were determined by requiring FDR < 0.05 between any two tissues. The same pipeline was also applied to the public data of PANDORA-seq [49] and CPA-seq [50] to calculate the total RPM of tsRNAs derived from each tRNA isodecoder.

RiboTag raw reads were first mapped to rRNA reference sequences using Bowtie2 (v2.3.5). Reads that were mapped to rRNAs were discarded. The remaining reads were then mapped to the mouse genome (mm10) using STAR (v2.7.5). Only uniquely mapped reads were considered for further analysis. Gene expressions were calculated using the StringTie v1.3.5.

### Metric definition

Codon index was designed to measure the usage of the codon, i, calculated as follows:

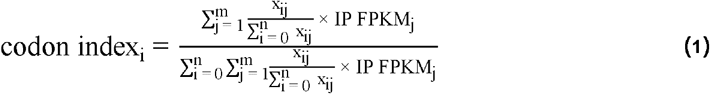

Here, x_ij_ denotes the number of occurrences of codon i in the gene j and FPKM_j_ denotes the FPKM value of gene j.

RSCU as defined by Sharp et al. [32] was calculated for each codon, j, as follows:

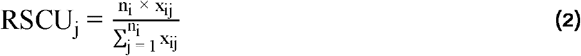

Here, n_i_ denotes the number of the synonymous codon for amino acid i, xij denotes the number of occurrences of codon j.

RSAU was calculated for each anticodon, j, as follows:

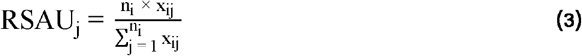

Here, ni denotes the number of the anticodon for amino acid i, x_ij_ denotes the number of occurrences of anticodon j.

RSIU was calculated for each Isodecoder, j, as follows:

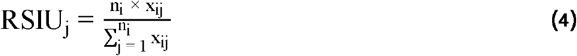

Here, n_i_ denotes the number of the isodecoders for anticodon i, x_ij_ denotes the number of occurrences of isodecoder j.

Amino acid composition was calculated for each amino acid, i, as follows:

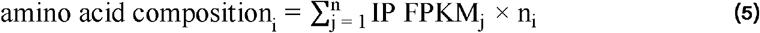

Here, n_i_ denotes the number of codons encoding amino acid i for gene j, IP FPKM_i_ denotes the FPKM value of gene j in IP of RiboTag.

### tRNA and translatome analyses

Permutation was performed by randomly switching the anticodons of the isodecoders and regrouping them into anticodons according to the permutated anticodons. We compared the mean CVs of anticodon expression, RSAU values, and tRNA isotype expression with 10,000 times permutations. The *P* values of permutation analyses were determined by calculating the fraction of permutations with the above values greater (isoacceptor expression, isotype expression) or less (RSAU) than the observed data.

Translational efficiencies were calculated as the ratio between the FPKMs of IP and the input of RiboTag. Only the genes with FPKM > 1 in both Input and IP samples were used in the downstream analyses. RSCU values were calculated as previously described by Sharp et al. [32] based on highly translated genes (top 5% translational efficiency). The coding region of the longest coding isoform of each gene was used for codon analyses. For comparison, RSCU values based on randomly sampled 5% genes with FPKM > 1 in both Input and IP samples were also calculated. Significance was determined by Wilcoxon signed-rank test.

tAI was calculated by R package tAI [36]. tAIs using different tRNA pools were calculated for highly translated genes (top 5% TE), medium translated genes (medium 5% TE), and low translated genes (bottom 5% TE). The significance between them was based on Wilcoxon signed-rank test. Data visualization and plotting were performed using ggplot2, ggrepel, and ggforce R packages.

The correlation analyses between isoacceptor abundances and the codon compositions of the top 5% highly translated genes of each tissue were based on the general codon-anticodon recognition rules for tRNA genes [36]. Codons recognized by multiple anticodons as well as anticodons that recognize multiple codons were repeated to form one-to-one codon-anticodon pairs.

## Ethical statement

Animal experiments were licensed with the approval No. SYSU-IACUC-2021-000089 and performed in agreement with the guidelines of the Animal Research Committee of the First Affiliated Hospital, Sun Yat-sen University.

## Data availability

The raw sequencing data of DM-tRNA-seq and RiboTag in this study have been deposited in the Genome Sequence Archive (GSA) at the National Genomics Data Center (https://bigd.big.ac.cn/) [54,55], Beijing Institute of Genomics, Chinese Academy of Sciences, and China National Center for Bioinformation (CNCB) (GSA: CRA005907 with BioProject: PRJCA008001; reviewer accessible link: https://ngdc.cncb.ac.cn/gsa/s/o8951Ujz), which are accessible at https://ngdc.cncb.ac.cn/gsa.

## CRediT author statement

**Peng Yu:** Methodology, Validation, Visualization, Writing – original draft. **Siting Zhou:** Software, Formal analysis, Visualization, Writing – original draft. **Yan Gao:** Investigation, Funding acquisition. **Yu Liang:** Investigation. **Wenbing Guo:** Formal analysis. **Dan Ohtan Wang:** Writing - Review & Editing. **Shuaiwen Ding**: Writing - Review & Editing. **Shuibin Lin:** Conceptualization, Writing – review & editing, Supervision, Project administration, Funding acquisition. **Jinkai Wang:** Conceptualization, Writing – review & editing, Supervision, Project administration, Funding acquisition. **Yixian Cun:** Conceptualization, Writing – review & editing, Supervision, Project administration. All authors read and approved the final manuscript.

## Competing interests

The authors have declared no competing interests.

## Acknowledgments

We are grateful to Professor Jianrong Yang for his constructive discussion and comments. This work was supported by the National Key R&D Program of China (Grant No.2018YFA0107200) to JW, the National Natural Science Foundation of China (Grant Nos.31970594 to JW; Grant Nos.81922052 and 81974435 to SL; Grant Nos.31971335 to D.O.W.), the Natural Science Foundation of Guangdong, China (Grant No.2019B151502011 to SL; 2021A1515110650 to YG), China Postdoctoral Science Foundation (Grant No. 2021M703755) to YG.

## Supplementary material

**Figure S1.**
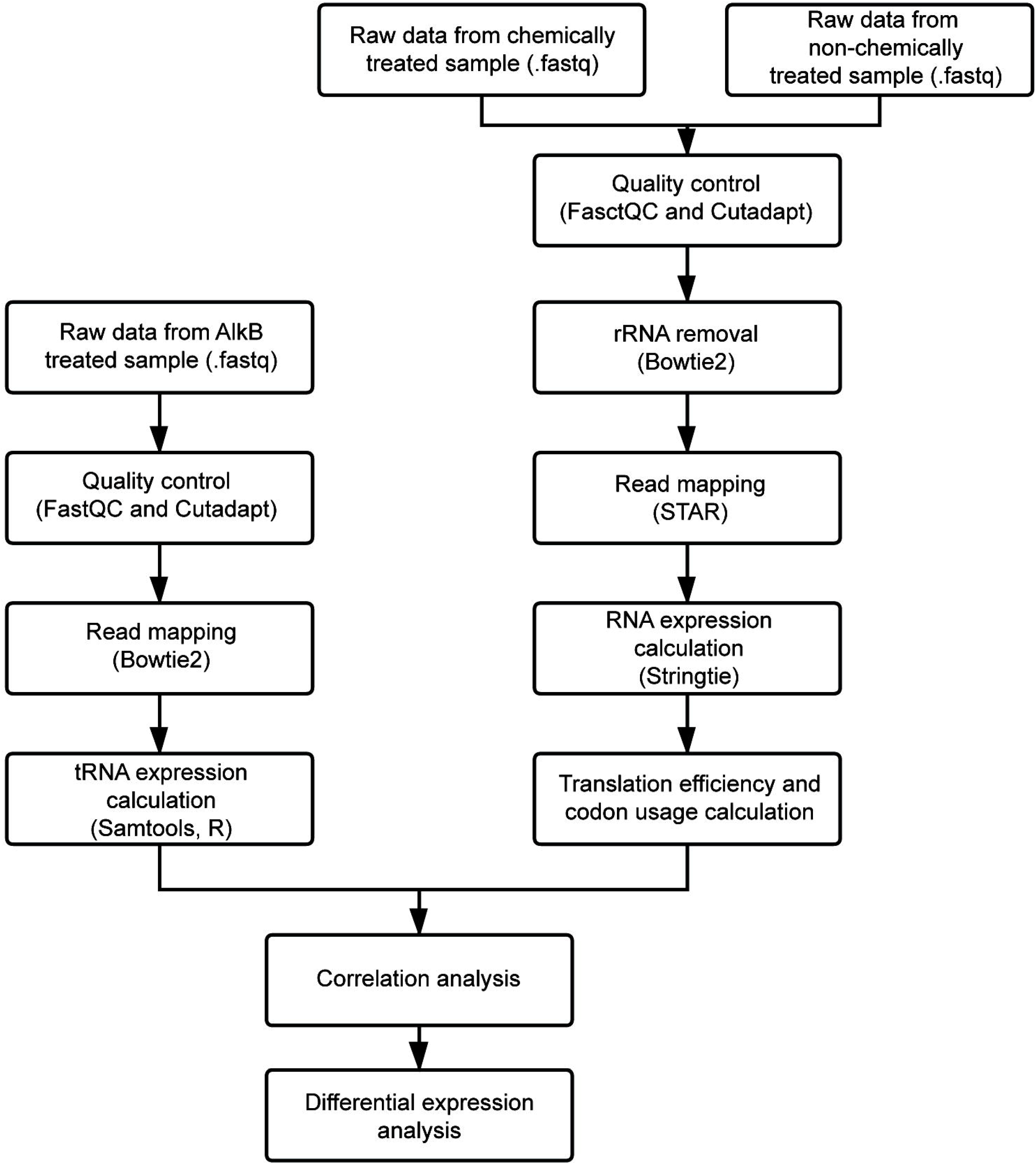
The bioinformatics analysis flow chart. The left flow chart shows the bioinformatic analysis steps of DM-tRNA-seq data; the right flow chart shows the bioinformatic analysis steps of RiboTag data.

**Figure S2.**
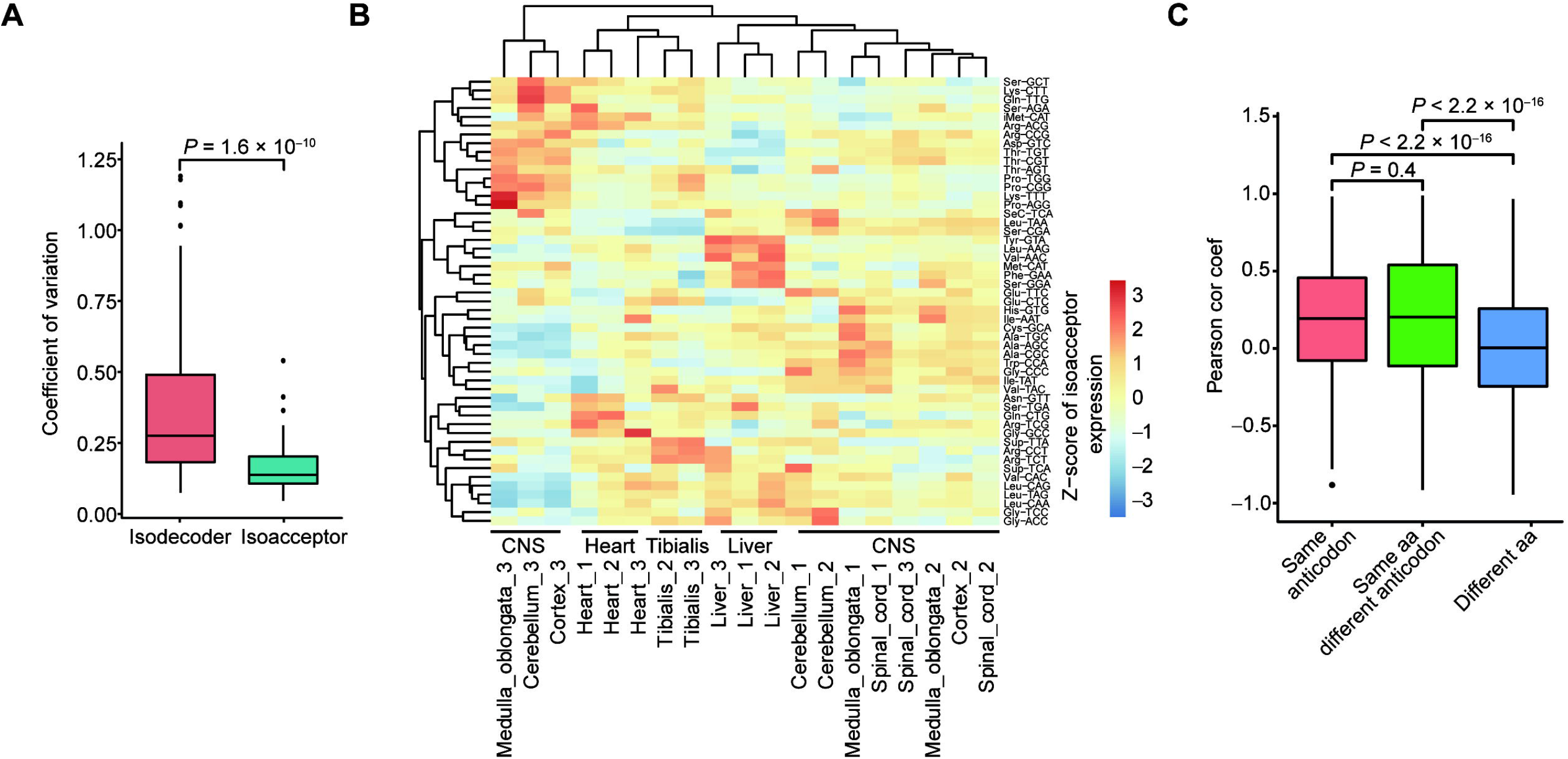
Dynamic expression of anticodons using published QuantM-tRNA seq data. **A.** Comparison of the coefficient of variation of isodecoders and isoacceptors among seven tissues, *P*-values of two-tailed Wilcoxon tests are indicated. **B.** Heatmap representing the Z-score of tRNA reads which collapsed by known isoacceptor groups in seven tissues. Two outliers were removed from the analysis (Cortex_1, Tibialis_1). **C.** Boxplot comparing the pairwise Pearson correlation coefficients of three groups of isodecoders: “same anticodon”, “same amino acid but different anticodon”, and “different amino acid” according to the corresponding anticodons and amino acids of the pairs of two isodecoders. CNS, central nervous system; aa, amino acid; Cor coef, correlation coefficient.

**Figure S3.**
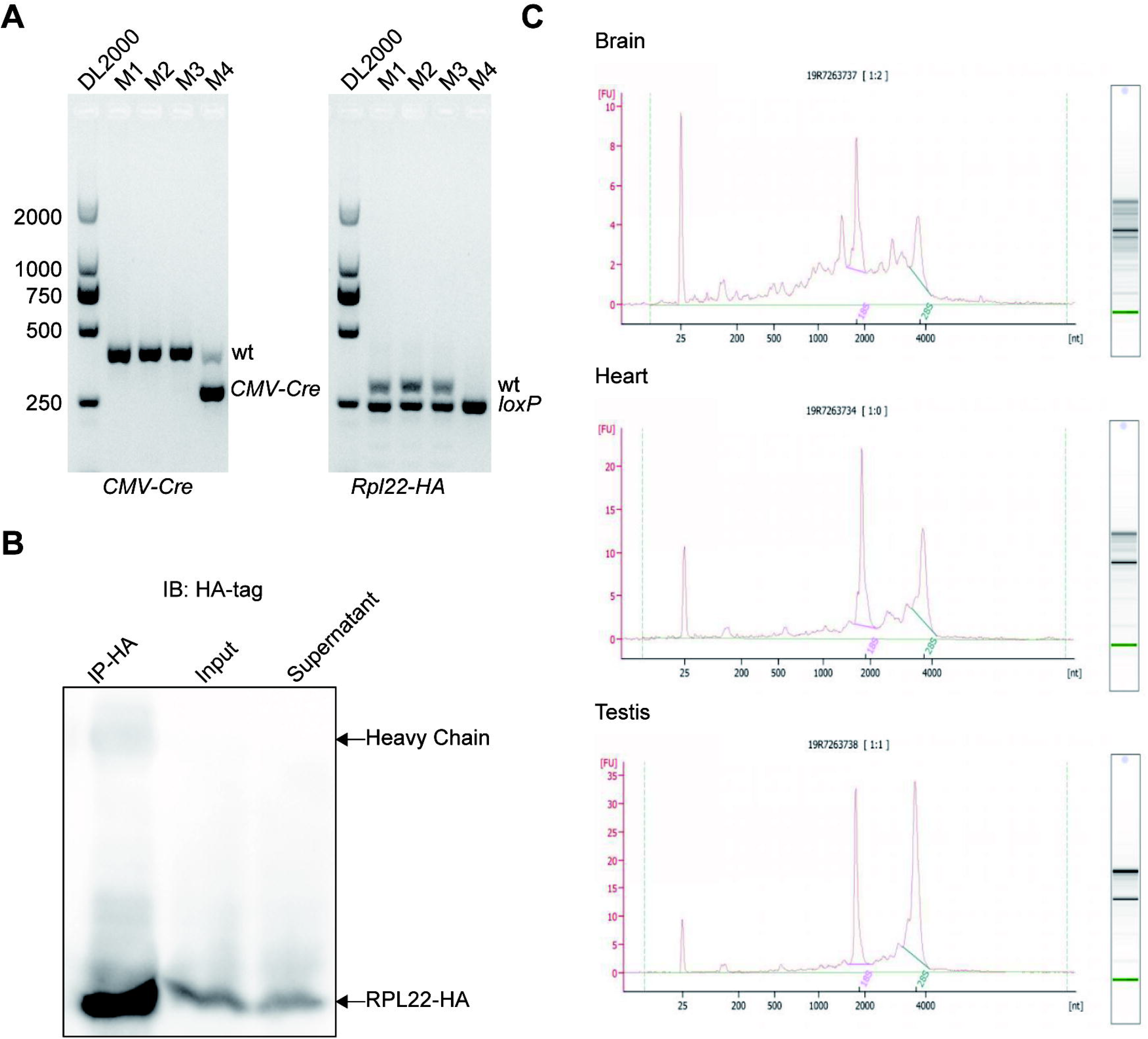
RiboTag analysis of translatomes in multiple mouse tissues reveals tissue-specific translational efficiency and codon usage biases. **A.** PCR products using primers that amplify CMV-Cre recombinase and the loxP-containing intron sequence of the *Rpl22* gene. The wild-type PCR product is 260 bp, while the mutant PCR product is 290 bp. **B.** Western blots using an anti-HA antibody demonstrate the presence of RPL22-HA specifically in anti-HA pellets versus supernatant. **C.** Agilent Technologies 2100 Bioanalyzer electropherogram analysis of total RNA from brain, heart, and testis immunoprecipitates. M, mouse No.; wt, wild type; IP, immunoprecipitation.

**Figure S4.**
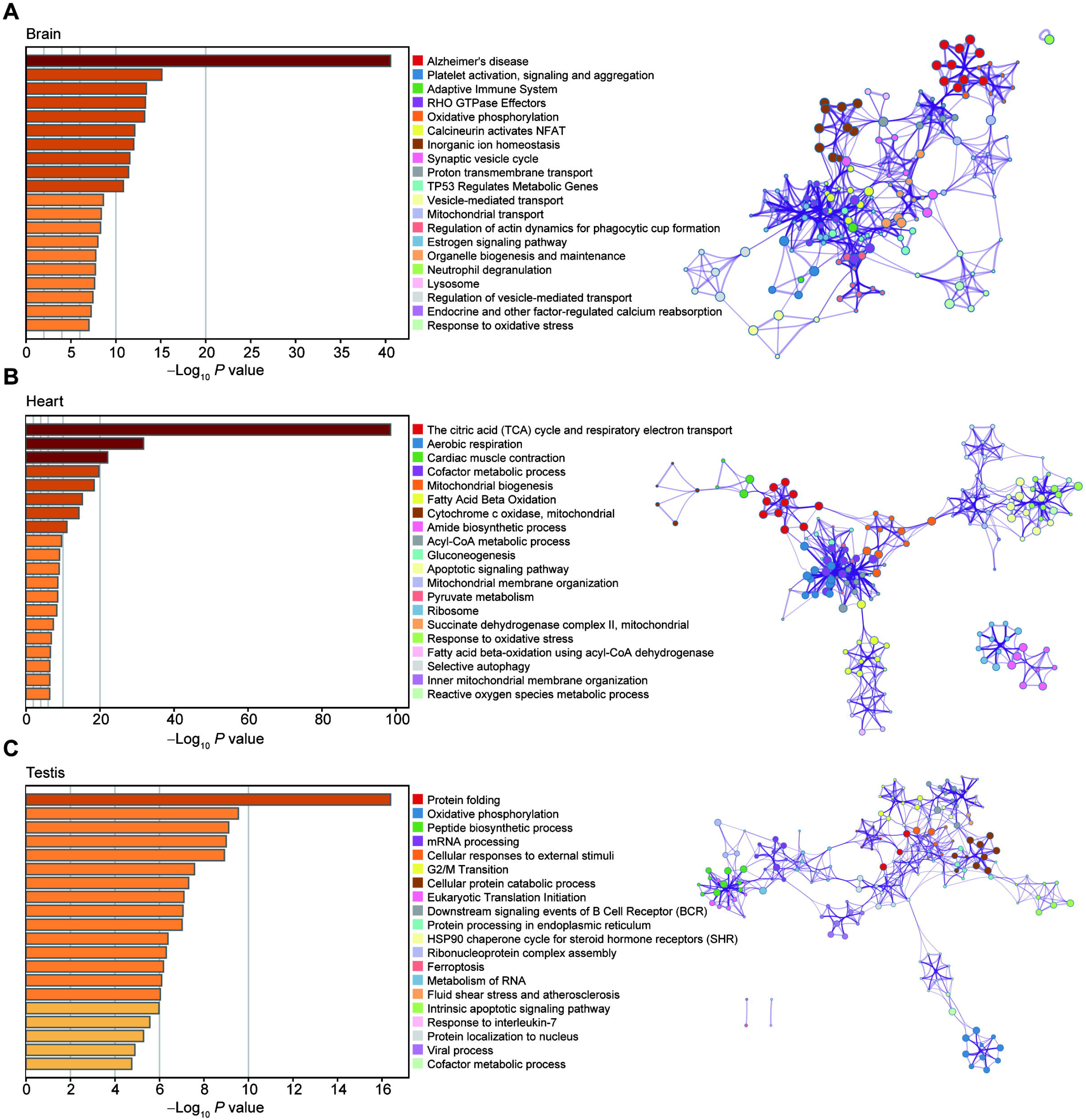
Metascape enrichment analysis of highly translated genes in different tissues. **A.–C.** Metascape enrichment analysis of top 5% highly translated genes in brain (A), heart (B), and testis (C).

**Figure S5.**
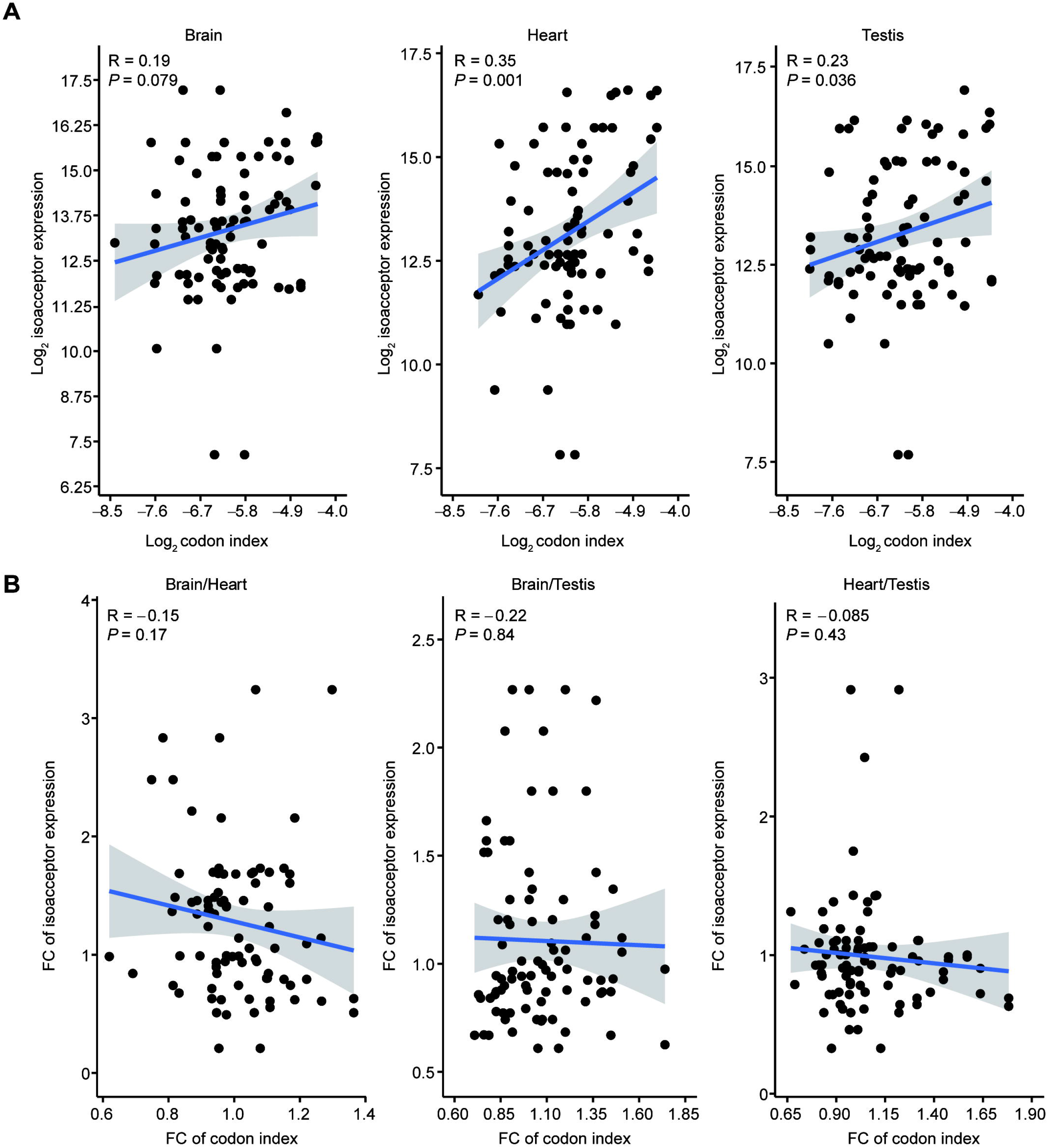
Pairwise correlation analyses of isoacceptor abundances and the codon compositions among the three tissues. **A.** Correlation analysis between isoacceptor abundances and the codon compositions. **B.** Correlation analysis between the fold change of isoacceptor abundances and the fold change of codon compositions. FC, fold change.

**Figure S6.**
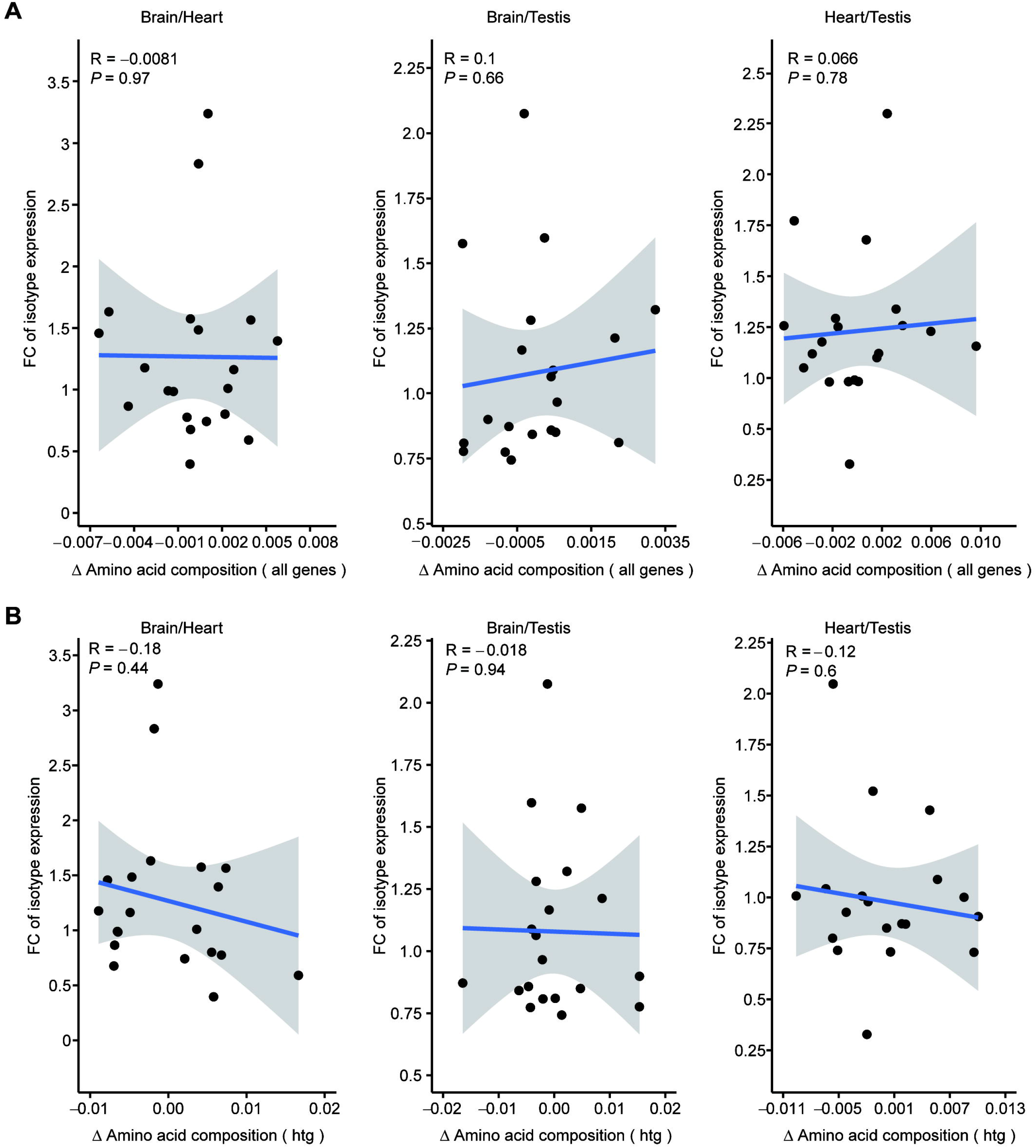
Pairwise correlation analyses of tRNA isotype expression and amino acid compositions among the three tissues. **A.** and **B.** Pairwise correlation analyses of the three tissues between delta amino acid composition and the fold change of tRNA isotype expression of all genes (A) and highly translated genes (B) respectively. htg: highly translated genes.

**Figure S7.**
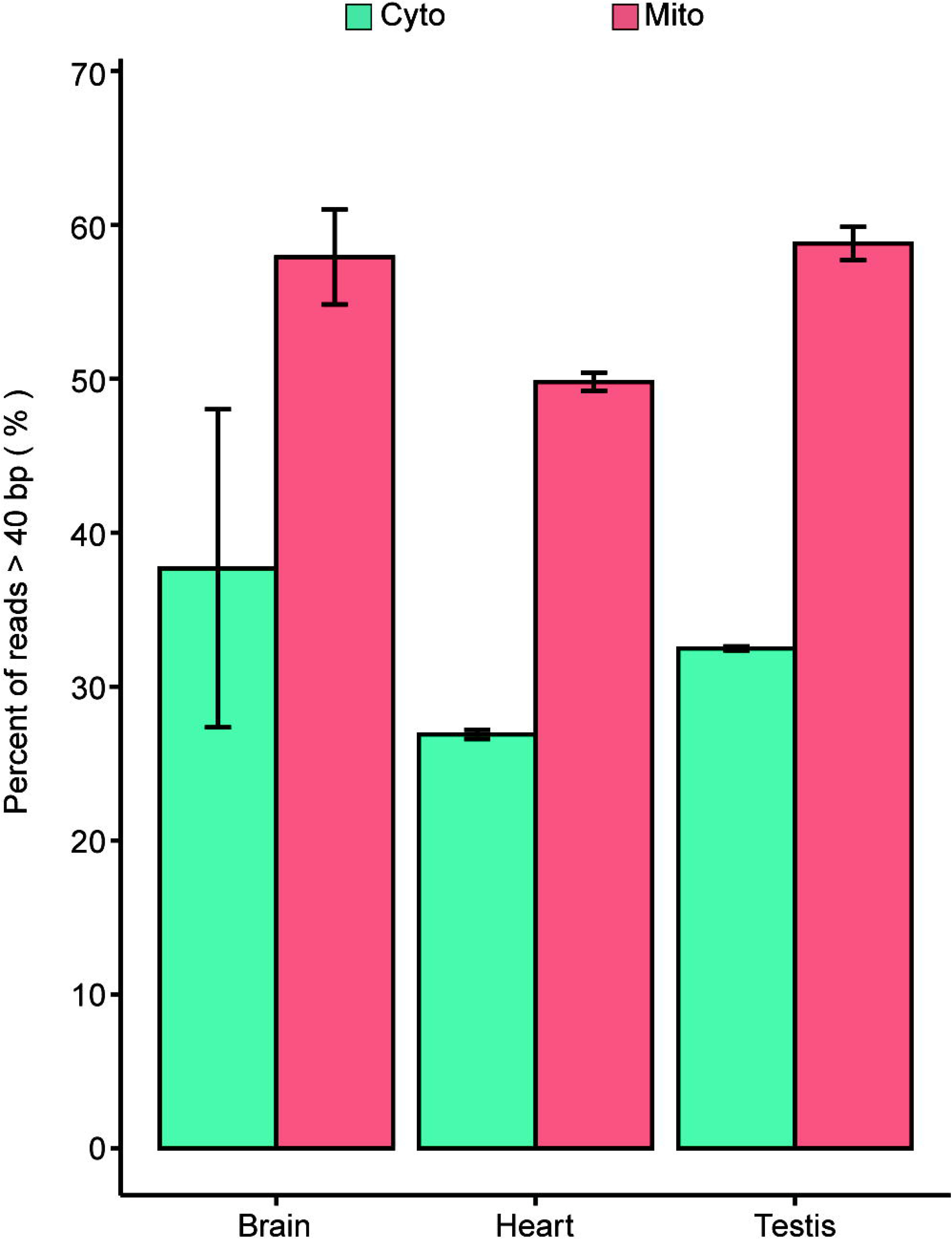
Comparison of the percentages of reads with length greater than 40 bp of cytosolic and mitochondrial tRNA. Cyto cytosolic; Mito, mitochondrial.

**Figure S8.**
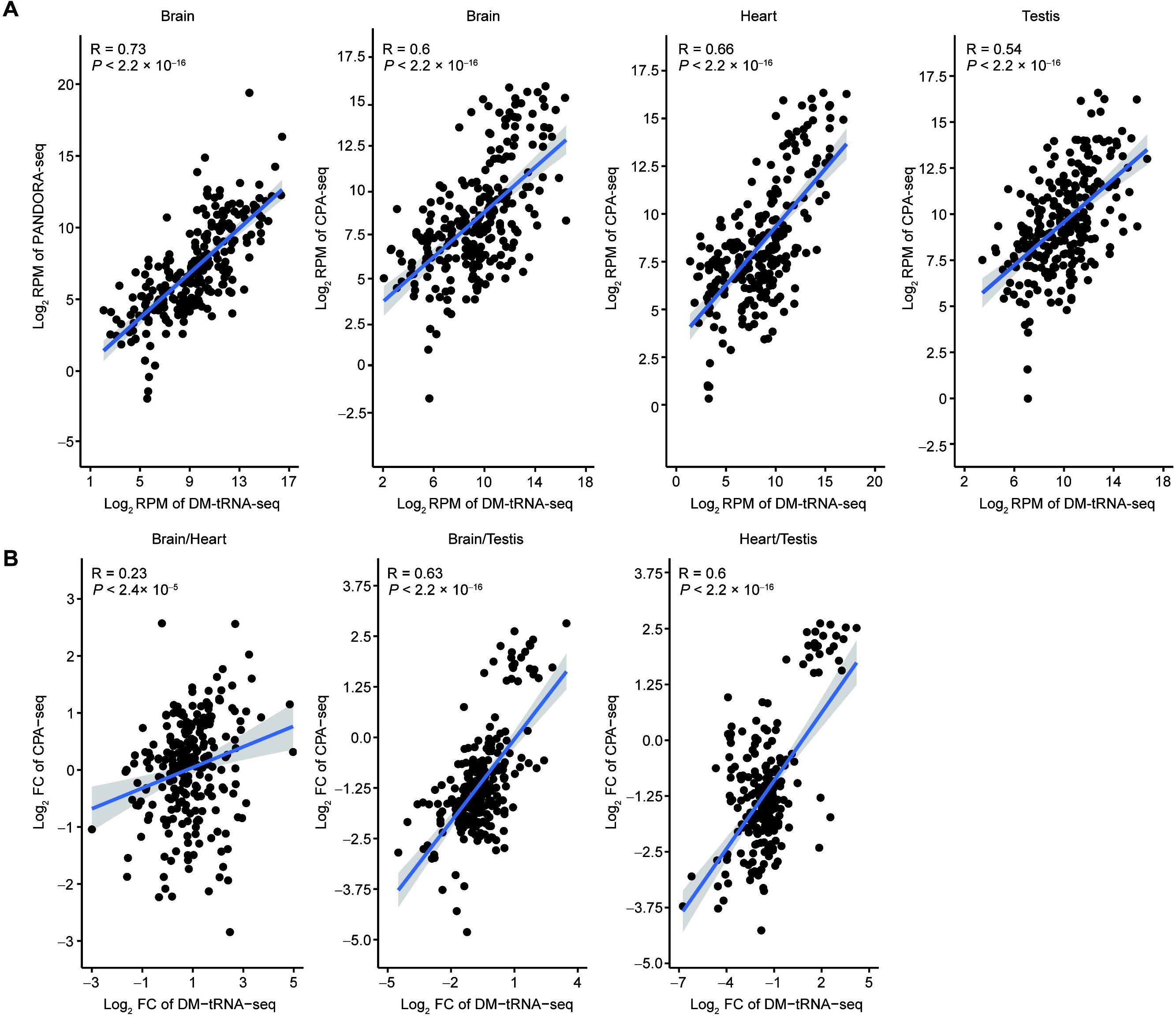
Correlation analyses of tRNA and tsRNA among the three tissues. **A.** Correlation analysis between tsRNA and tRNA on isodecoder level. **B.** Correlation analysis between the fold change of tsRNA and the fold change of tRNA.

**Table S1 The counts and expression of tRNAs**

**Table S2 The gene FPKMs of IP and Input of RiboTag**

